# Non Parametric Differential Network Analysis for Biological Data

**DOI:** 10.1101/2023.12.08.570801

**Authors:** Pietro Hiram Guzzi, Arkaprava Roy, Pierangelo Veltri

## Abstract

Rewiring of molecular interactions under different conditions causes different phenotypic responses. Differential Network Analysis (also indicated as DNA) aims to investigate the rewiring of gene and protein networks. DNA algorithms combine statistical learning and graph theory to explore the changes in the interaction patterns starting from experimental observation. Despite there exist many methods to model rewiring in networks, we propose to use age and gender factors to guide rewiring algorithms. We present a novel differential network analysis method that consider the differential expression of genes by means of sex and gender attributes. We hypothesise that the expression of genes may be represented by using a non-gaussian process. We quantify changes in nonparametric correlations between gene pairs and changes in expression levels for individual genes. We apply our method to identify the differential networks between males and females in public expression datasets related to mellitus diabetes in liver tissue. Results show that this method can find biologically relevant differential networks.

## 1 Introduction

The always more performing high-throughput technologies for studying genes, proteins and non-coding RNA allowed us to identify and study many associations among changes in their abundance and diseases [12]. The availability of data has also motivated the introduction of novel methods of analysis based on a systemic perspective by using network science. Networks have been used to represent the set of associations among biological molecules in a given state starting from experimental observation [9]. An interesting application of network modelling is the possibility of gathering information by comparing biological networks associated with different conditions (e.g. healthy vs diseases). Differential Network Analysis (DNA) has been introduced to model differences between two conditions represented as two distinct networks in a single network (differential network) which model the differences [13, 8, 10, 5]. DNA methods have been used to compare experimental configurations (e.g. two different drugs) or phenotypes. Recently, many observations have evidenced that the age and sex of patients may have a role in causing different responses to drugs, outcomes of diseases and incidence of comorbidities for many complex chronic diseases [25, 11, 24]. Experimental observations evidenced that both the incidence and progression of some diseases have remarkable differences considering sex and age as factors. For instance, patients affected by diabetes are more likely to develop comorbidities as they grow older, as well as studies about mortality caused by COVID-19 pandemia, showed higher number of deaths in older males wrt female [3, 17].Consequently, the necessity of defining novel algorithms to identify motivations or possible candidates that causes such differences at the molecular level and depending on age or genders, arises.

DNA algorithms aim to identify changes in measures of association in terms of network rewiring, which can distinguish different biological conditions *𝒞*_∞_, *𝒞*_∈_. Let *𝒞*_∞_, *𝒞*_∈_ be two different biological conditions in two different networks representing molecular interactions, DNA algorithms aim to identify changes in network rewiring (and differences) among the two networks that may be possible candidates associated to changes among *C*_1_ and *C*_2_. Given two gene expression datasets corresponding to two experimental conditions, DNA algorithms first derive two networks *𝒩*_∞_, *𝒩*_∈_ corresponding to the examined condition. Each network usually has a node for each gene of the dataset, while a weighted edge between nodes represents the association or the casual dependency among them and the weight represents the strength of the association. Finally, a single network _⌈_ describing the changes is calculated. The final network represents the rewiring of association in condition, and it has been used in the past to study changes associated with pathological conditions [11, 2]. Given two datasets corresponding to gene expressions related to two different biological experiments, DNA is based on the following steps. Firstly two networks *𝒩*_∞_, *𝒩*_∈_ corresponding to the examined conditions are defined. Each node in a network corresponds to a gene in the dataset, and weighted edges between nodes represent the association or the casual dependency among genes, while the weight on an edge represents the strength of the associations among genes. Then, DNA algorithmic implementations compute a new single network *𝒩*_⌈_ describing the changes (i.e. the differents among networks *𝒩*_∞_ and *𝒩*_∈_). The resulting network represents the rewired associations that can be used to study biological triggers causing rewiring differences. This may be used to study variations associated with pathological conditions as reported in [11, 2].

Many existing methods are based on the hypothesis that experimental observation such as gene expression values come from parametric distribution, e.g. normal, paranormal, binomial or Poisson distribution [16, 6, 18, 12]. NGS data, different to microarray technology, are similar to count data, thus parametric distribution hypothesis sometimes does not hold. Consequently, there is a need for the introduction of non-parametric methods for DNA analysis.

We here propose a novel DNA method to identify differential edges among two networks and integrate differential expressions between nodes (i.e. genes) also using gender-based differences [5]. Moreover, gene expression level is statistically predicted by using multivariate count data, and the conditional dependence graph is built by using pairwise Markov random fields [21]. Differently to existing methods where gene expression values are obtained by using parametric distribution (such as normal, paranormal, binomial or Poisson) [16, 6, 18, 12], DNA analysis requires non-parametric methods. In a nutshell, the proposed DNA algorithm works as follows. We build two graphs for the two tested conditions; then, we derive the final graph from the previous two. Finally, we prune the resulting graph by admitting only edges incident to at least one differential expressed gene. The proposed DNA has been used with genes dataset related to patients affected by diabetes and we use results to identify the differential networks also using gender attributes (i.e. male and female patients).

The paper is structured as follows: Section 2 discusses state-of-the-art methods; Section 3 presents the proposed approach; Section 4 discusses the results of some experiments; finally Section 5 concludes the paper.

## 2 Related Work

DNA is largely used to identify differentially expressed genes between groups of samples, and thus useful to compare genes from patients with a particular disease compared to healthy subjects. In molecular biology and bioinformatics it is used for identifying genes that are differentially expressed between groups of samples, such as those from patients with a particular disease compared to healthy individuals.

DNA algorithms aim to identify changes in the network structures between two states, or conditions [23]. In biology, DNA algorithms have been used to identify changes between the healthy and diseased status of the same biological system [9]. There exist some different formulations of the problem, we here focus on networks with the same node sets and different edge sets. Formally, given two different conditions *𝒞*_1_, and *𝒞*_2_, represented by means of two graphs *G*_1_(*V, E*_1_) and *G*_1_(*V, E*_2_), DNA aims to identify changes between them.

When dealing with biological systems it should be noted, that nodes are directly measurable, while edges among them should be derived from a set of observations over time. For instance, when considering gene networks derived from microarray experiments, nodes are fixed while edges should be inferred from the observations by means of *statistical graphical models* [14, 22, 7]. In a statistical graphical model, we use a graph *G* = (*V, E*), and each node *v* ∈ *V* is associated with a set of *m* random variables *X*_1_, …, *X*_*m*_ representing quantitative measurements of *v*, and edges are inferred from *X*_1_, …, *X*_*m*_. We focus on **undirected** graphs. In such models differential associations are measured by analysing the the difference of partial correlations between experimental data of two conditions. Changes are measured by means of specific statistics test defined to measure the modification among correlation between entities. Moreover, the changes in gene expression levels are quantified by using the classical Student’s t-test statistics [27]. Then, the two test statistics are integrated into a single optimisation model which aims to evidence the hierarchical structures of networks. Nevertheless, some hypotheses of the previous models (e.g. gaussian distribution of data) are not valid in all the experimental conditions, therefore non-parametric methods have been introduced. These methods are in general computationally efficient and often easier to implement and the results can be more interpretable. The main limitation of these methods is that they require the data to adhere to specific distributional assumptions, and if these assumptions are violated, the results can be biased or incorrect.

Some works considered a nonparanormal distribution of data (or Gaussian copula) instead of normality or multivariate normality of data [1] and they used a rank-based correlation matrix, such as Spearman correlation or Kendal’s *τ*. Since the nonparanormal model presents some restrictions on the nature of data, some conditionally-specified additive graphical models have been proposed such as graphical random forest and kernel-based estimators [23].

In particular such models have been used for brain data and counts based data, such as sequencing. To overcome the time limitations of the non-parametric methods, efficient Bayesian models have been proposed [23]. Such methods are based on the calculation of probabilities of the edges among data by inferring their likelihood. Some of the proposed methods used different heuristics to infer such probabilities which are hard to derive from data, such as in [21]. We here selected this last method which outperforms the other state-of-the-art methods.

Non-parametric methods make fewer assumptions about the underlying data distribution and they are based on data-driven techniques to assess the differences in network connectivity between conditions. These methods are more flexible and robust, as they do not assume a specific data distribution and can handle complex and non-linear relationships between nodes in the network. On the other hand, they can be computationally intensive and less interpretable.

Choosing between parametric and non-parametric differential network analysis depends on the nature of the data, the underlying assumptions, and the research question. Researchers often perform sensitivity analyses and cross-validate their results to ensure the robustness and reliability of the findings. Additionally, combining information from both approaches may provide a more comprehensive understanding of the differential network structure.

## 3 The proposed pipeline

This section explains the method we designed and implemented as depicted in Figure 1. The method starts by gathering expression data grouped by tissues and for each tissue we build two different datasets filtered by using gender of the individual they belongs to.

**Fig. 1.**
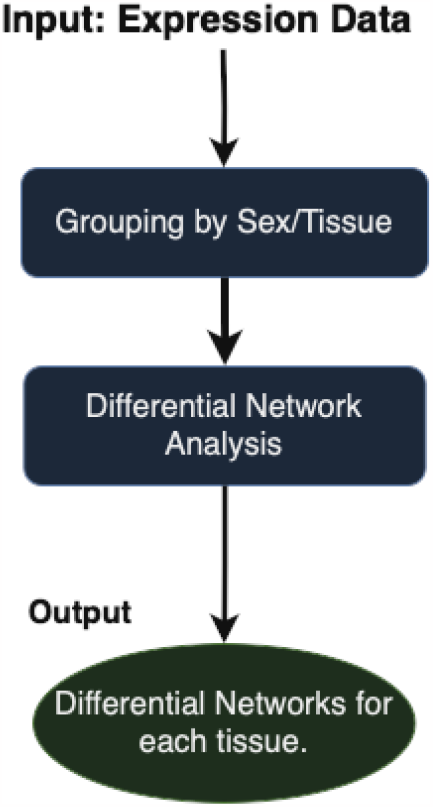
Figure depicts the main steps of the experiments we performed.

### 3.1 Non Parametric Differential Network Analysis

Let us suppose that two biological conditions *𝒞*_1_ *𝒞*_2_, have been investigated by means of gene expression analysis giving two different expression datasets encoded in two matrices *N*×*M* (N samples, M genes) *X*^1^, *X*^2^. Each row of *X*^*j*^ stores the expression values of *M* genes of the *i* sample. Therefore 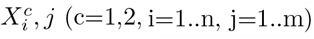 denotes a pair of *N* × *M* matrices.

Many approaches suppose that gene expression datasets are samples from two multivariate normal distributions. We here do not hold this hypothesis, so we hypothesise that two gene expression datasets are samples from non-parametric distributions.

Let 𝒫_1,2_ (∥*P* ∥ = *nxn*) be two matrices representing the relation among nodes. Both matrices represent the conditional independence among nodes [21].

We may define the differential matrix between two conditions as the difference as 𝒫_1_ and 𝒫_2_.

Following pair-wise MRF [26, 4], we consider the following joint probability mass function for *P* -dimensional count-valued data *X* as in [21],

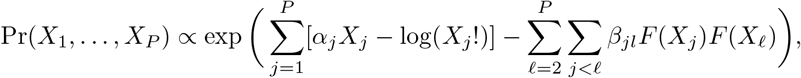

where *F* (·) is a monotone increasing bounded function with support [0,∞). We let *F* (·) = (tan^−1^( ·))^*θ*^ for some positive *θ* ∈ ℝ^+^ to define a flexible class of monotone increasing bounded functions. The exponent *θ* provides additional flexibility, including impacting the range of *F* (*X*), 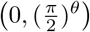. The parameter *θ* can be estimated along with the other parameters, including the baseline parameters *α* controlling the marginal count distributions and the coefficients *β*_*jl*_ controlling the graphical dependence structure. For simplicity and interpretability, we estimate *θ* to minimize the difference in covariance between *F* (*X*) and *X* following [21] For detailed descriptions of the method, readers are encouraged to check [21].

If *β*_*jℓ*_ = 0, we have *X*_*j*_ and *X*_*ℓ*_ to be conditionally independent i.e. *P* (*X*_*j*_, *X*_*ℓ*_ | *X*_−(*j,ℓ*)_) = *P* (*X*_*j*_ | *X*_−(*j,ℓ*)_)*P* (*X*_*ℓ*_ | *X*_−(*j,ℓ*)_), where *X*_−(*j,ℓ*)_ stands for all the variables excluding *X*_*j*_ and *X*_*ℓ*_. Our estimated graphical relation would rely on 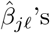, and thus, our model is a probabilistic model that encodes the conditional independence structure in a graph.

Consequently, we define a differential network as the difference 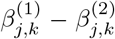 for each edge (j,k) where 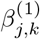 and 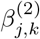are estimated coefficients under two conditions 1 and 2. From the MCMC samples, we can get the posterior estimates of these differences as 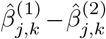. Alternatively, we may be able to compute other posterior summaries such as 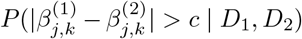, which is the posterior probability that 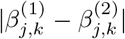 is greater than some pre-specified cutoff given the two datasets, denoted as *D*_1_ and *D*_2_.

## 4 Experimental Results

Figure 1 depicts the main steps of our experiments. We start by considering GTEx gene-expression data [15]. Genotype-Tissue Expression (GTEx) data portal [15], which is a publicly available resource containing expression data of patients integrated with information related to the tissue of provenance, sex and age (grouped into six classes). The current version of the GTEx database accessed on February 01st stores 17382 samples of 54 tissues of 948 donors, see at https://gtexportal.org/home/tissueSummaryPage [17, 20, 19]. We downloaded data and integrated it with information related to sex and age with an ad-hoc realised script. Then for each tissue, we split data into male and female samples. For each tissue, we randomly selected the same number of samples.

We used Conga [21] R package to derive non-parametric differential networks. Conga receives as input two datasets corresponding to the different experimental conditions. We ran conga using default parameters. We here show results for genes related to diabetes expressed in the liver tissue in Figure 2.

**Fig. 2.**
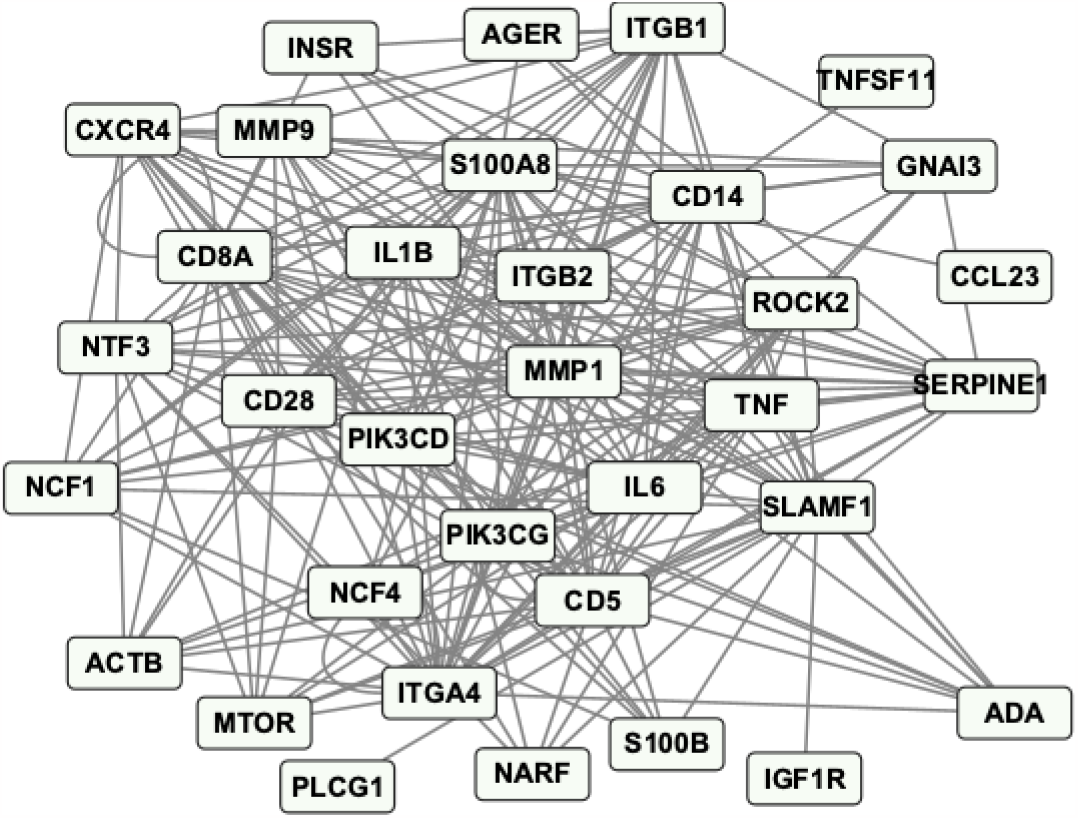
Figure represents a differential network between males and females in liver tissue.

To explain this network’s biological and medical relevance, we perform a Gene Ontology analysis using the String Database as depicted in Figure 3.

**Fig. 3.**
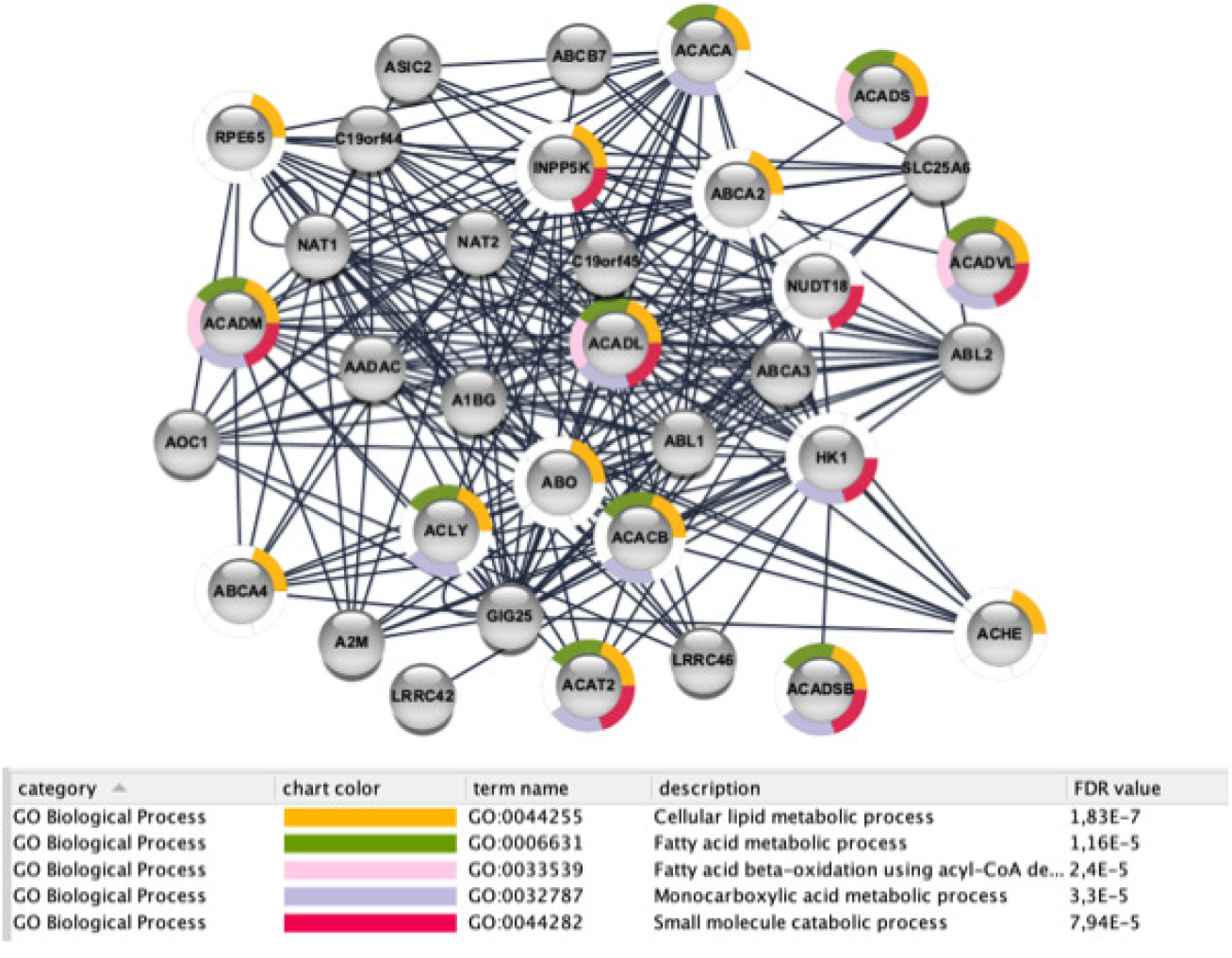
Figure depicts the main enriched function of the obtained network. We used the String Database and we selected Gene Ontology Biological Process. All the functions are enriched with a p-value after false discovery rate correction value less than 0.01

## 5 Conclusion

The results presented in the article demonstrate the effectiveness of the proposed non-parametric approach for Differential Network Analysis (DNA) in uncovering meaningful biological insights from gene expression data. The analysis focuses on gender-based differences in liver tissue using the Genotype-Tissue Expression (GTEx) database.

The differential network provides a visual representation of how gene interactions change between males and females, offering insights into gender-specific molecular relationships. To understand the biological relevance of the identified differential associations, we performed a Gene Ontology analysis using the STRING Database. The enriched Gene Ontology terms shed light on the biological processes that may be influenced by gender-specific gene interactions in liver tissue.

The results highlight specific genes and interactions that exhibit gender-based differences. This information is valuable for understanding why certain diseases or conditions might affect males and females differently. By identifying gender-specific molecular associations, researchers can potentially uncover molecular mechanisms underlying sex-related disparities in disease susceptibility, progression, or response to treatments.

The success of the approach in identifying known gender-specific molecular differences validates the effectiveness of the method. If the identified differential network aligns with existing knowledge about gender-related gene interactions or processes, it strengthens the credibility of the approach and its ability to reveal biologically relevant insights.

While the results provided promising insights, the analysis may also be influenced by factors like data quality, sample size, and the choice of statistical parameters. Additionally, the results might be specific to the dataset used (GTEx), and their generalizability to other datasets or tissues should be considered.

Understanding gender-specific molecular differences has implications for personalized medicine. The identified genes and interactions could potentially serve as biomarkers for predicting disease risk, prognosis, or treatment response based on an individual’s gender. This could lead to more tailored and effective medical interventions.

In summary, the results presented in the article showcase the potential of the non-parametric Differential Network Analysis method to uncover gender-based differences in gene interactions. The identified differential network and enriched Gene Ontology terms provide insights into the molecular underpinnings of gender-related disparities in liver tissue. This approach has broader implications for understanding sex-specific responses to diseases and treatments, advancing the field of personalized medicine.

## 6. Acknowledgements

Authors thank Francesca Cortese for the preliminary analysis conducted.

